# Responses to membrane potential-modulating ionic solutions measured by magnetic resonance imaging of cultured cells and *in vivo* rat cortex

**DOI:** 10.1101/2024.04.02.587661

**Authors:** Kyeongseon Min, Sungkwon Chung, Seung-Kyun Lee, Jongho Lee, Phan Tan Toi, Daehong Kim, Jung Seung Lee, Jang-Yeon Park

## Abstract

Membrane potential plays a crucial role in various cellular functions. However, existing techniques for measuring membrane potential are often invasive or have limited recording depth. In contrast, magnetic resonance imaging (MRI) offers noninvasive imaging with desirable spatial resolution over large areas. This study investigates the feasibility of utilizing MRI to detect responses of cultured cells and *in vivo* rat cortex to membrane potential-modulating ionic solutions by measuring magnetic resonance parameters. Our findings reveal that depolarizing (or hyperpolarizing) ionic solutions increase (or decrease) the *T*_2_ relaxation time, while the ratio of bound to free water proton shows the opposite trend. These findings also suggest a potential approach to noninvasively detect changes in membrane potential using MRI.

## Introduction

Membrane potential is a fundamental property of all living cells, influencing crucial cell functions^1^ such as neuronal and myocyte excitability, volume control, cell proliferation, and secretion. In the fields of neuroscience, membrane potential is of significant importance, as neural activities arise from the dynamic propagation of this electric potential. From a clinical perspective, deviations from normal membrane potential levels contribute to various diseases, including seizures^2^, arrhythmia^3^, and hypoglycemia^4^. Given its scientific and clinical importance, effective methods to detect changes in membrane potential have long been sought.

The intracellular recording technique using sharp glass electrodes is a commonly used method to detect changes in membrane potential^5,6^. It provides real-time absolute recordings of membrane potential. Optical imaging is another major approach to detect changes in membrane potential. Voltage-sensitive dyes enable voltage imaging at the cellular level^7^. Fluorescent calcium indicators detect calcium transients associated with neuronal activation^8^. Label-free optical imaging techniques^9^ utilize various contrast mechanisms such as cell membrane deformation, which are directly coupled with changes in membrane potential. Despite their efficacy, these methods have limitations when applied to intact biological systems due to their invasive nature, requiring procedures such as craniotomy.

As noninvasive techniques for directly or indirectly detecting brain activation *in vivo*, several imaging modalities have been developed, including electroencephalography (EEG)^10^, magnetoencephalography (MEG)^11^, and magnetic resonance imaging (MRI)^12^. EEG and MEG detect electric potentials on the scalp and extracranial magnetic fields, respectively, which are directly induced by neuronal activity in the brain. Although these techniques are noninvasive and provide excellent temporal resolution of milliseconds or less, they are constrained by shallow imaging depth and spatial localization challenges^13^.

In contrast, MRI enables noninvasive imaging with good spatial resolution of millimeters over a large brain volume, making it an appropriate tool for *in vivo* functional brain imaging. To date, the mainstream of functional MRI (fMRI) utilizes hemodynamic responses driven by brain activation, such as the blood oxygen level-dependent (BOLD) effect^14^. However, while the BOLD contrast mechanism reflects dynamic changes in neuronal activity through neurovascular coupling, it provides inherently indirect and relatively slow, hemodynamic-responsive information of brain function^15^. On the other hand, many studies have attempted to explore the possibility of using MRI to directly detect neuronal activity^16,17^. These studies utilized neuronal current-dependent signal phase shifts^18-20^ and magnitude decay^21-24^, the Lorentz effect^25^, high temporal resolution^26,27^, ghost artifacts^28^, or cell swelling^29,30^, while concerns about sensitivity exist^31-36^.

In this study, we investigated the possibility of using MRI to detect membrane potential changes induced by modulating ionic solutions. Specifically, we explored how *T*_2_ relaxation time and magnetization transfer (MT) correlate with membrane potential changes, both *in vitro* and *in vivo. In vitro* experiments utilized two homogeneous and electrically non-active cells, i.e., neuroblastoma (SH-SY5Y) and leukemia (Jurkat) cell lines, providing a controlled environment free from hemodynamic effects. This aided in accurately evaluating changes in MR parameters with membrane potential. *In vivo* experiments, conducted on a rat model with a craniotomy-exposed cortex, aimed to reproduce the *in vitro* findings.

## Results

### *In vitro* changes in MR parameters induced by membrane potential

A non-excitable neuroblastoma cell line, SH-SY5Y, was selected to investigate the relationship between MR parameters and membrane potential modulated by ionic solutions. After culturing SH-SY5Y cells, they were suspended in extracellular media with various potassium ion concentrations ([K^+^]), while maintaining constant osmolarity by adjusting [Na^+^]. As the control condition, [K^+^] = 4.2 mM was selected. For depolarization conditions, [K^+^] = 20, 40, and 80 mM were used. For hyperpolarization conditions, [K^+^] = 0.2 and 1 mM were used. The suspended cells were concentrated with centrifugation in an acrylic container and scanned in a 9.4 T preclinical MRI system. The imaging slice was positioned 0.5 mm below the pellet-extracellular media interface to ensure that signals were acquired from cell pellet. Under each condition, *T*_2_ and MT parameters such as pool size ratio (PSR) and magnetization transfer rate (*k*_*mf*_) were measured (Fig. 1). The PSR value represents the ratio of hydrogen protons in macromolecules and free water, and *k*_*mf*_ represents the magnetization transfer rate of hydrogen protons from macromolecules to free water. In addition, the membrane potential of SH-SY5Y cells in each condition was separately measured via patch clamp recording.

**Fig. 1.**
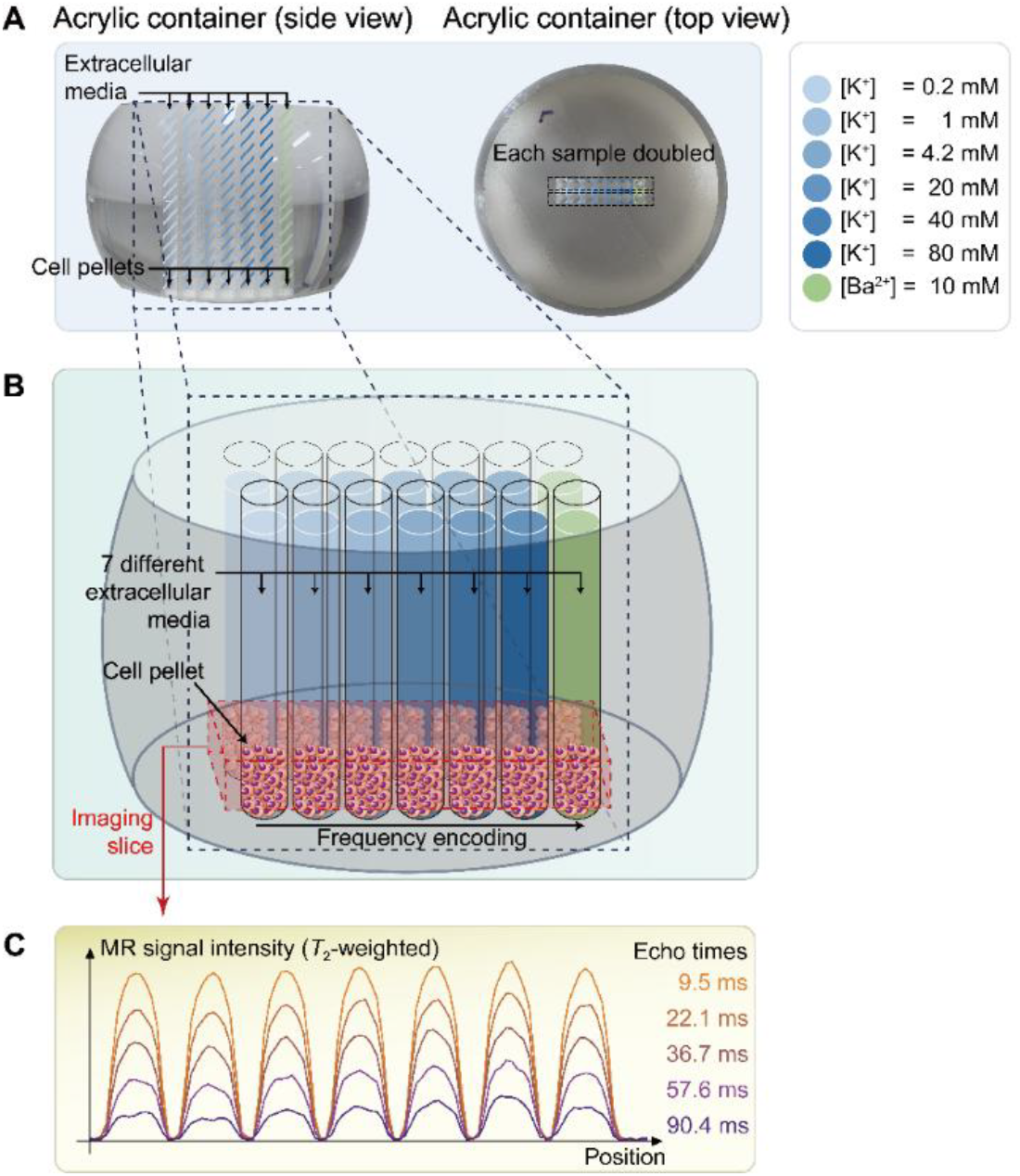
The schematic diagram of the *in vitro* experiment. (A) The picture on the left displays a side view of a double-sided cut spherical acrylic container with fabricated wells filled with extracellular media and cell pellets. As depicted in the top-view picture on the right, fourteen wells (matrix = 2 × 7) were created on the acrylic container, allowing each of the seven samples with six different K^+^ concentrations ([K^+^] = 0.2–80 mM) and one Ba^2+^ concentration ([Ba^2+^] = 10 mM) to be doubled in the same column for improved signal-to-noise ratio (SNR) in MR signal acquisition. (B) The image illustrates the configuration after loading cells into the wells and pelleting them at the bottom of the wells. The imaging slice was positioned 0.5 mm below the pellet-media interface to acquire signals predominantly from the cell pellets. (C) Representative one-dimensional *T*_2_-weighted MR signals with 5 selected echo times out of a total of 50 acquired echo times.

The changes of *T*_2_, PSR, and *k*_*mf*_ in SH-SY5Y cells when the membrane potential (*V*_*m*_) was modulated by varying [K^+^] are shown in Figure 2, alongside the actual *V*_*m*_ measured via patch clamp recordings. We conducted statistical analyses to assess the effect of changes in *V*_*m*_ from the control condition (Δ*V*_*m*_) on the MR parameters. A linear mixed-effect model was applied to account for inter-sample variability and repeated measurements. This model included MR parameters as dependent variables, Δ*V*_*m*_ as a fixed effect, and cell batch as a random effect. The analysis yielded the following relationships:

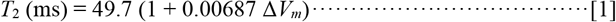

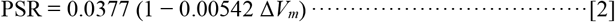

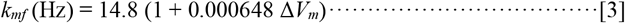

**Fig. 2.**
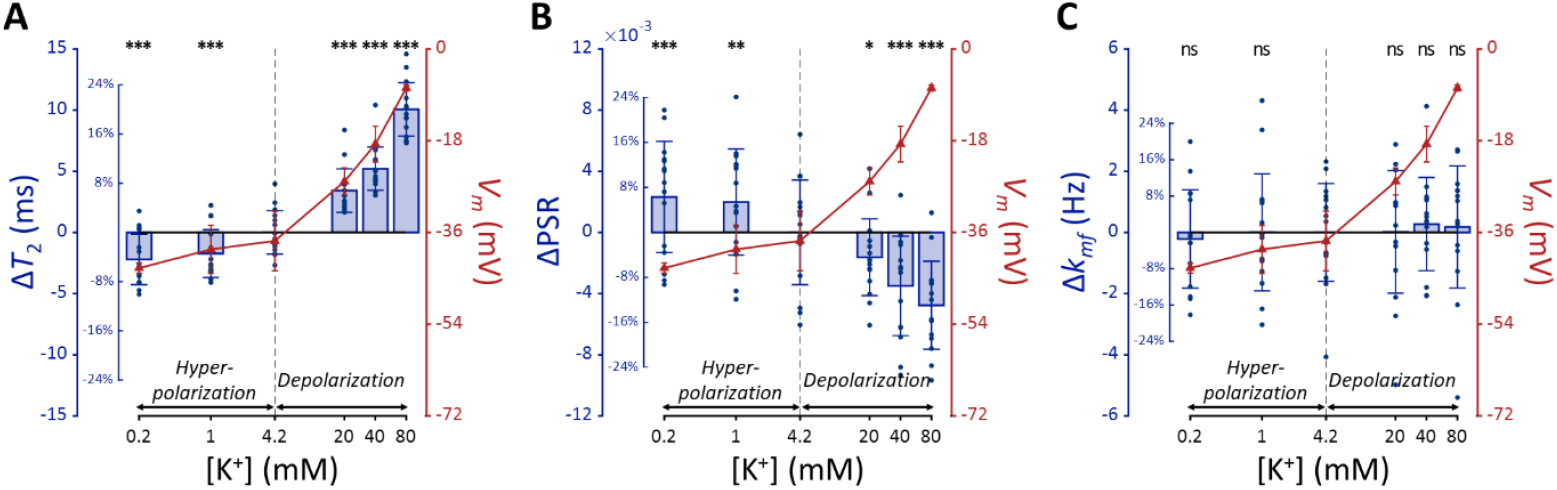
MR parameters and membrane potential (*V*_*m*_) of SH-SY5Y cells versus extracellular K^+^ concentrations ([K^+^]). Changes in (A) *T*_2_, (B) PSR, and (C) *k*_*mf*_ are displayed with blue bars (*n* = 15). Membrane potentials are plotted with red triangles (*n* = 3). The abscissa is in logarithmic scale. Error bars denote standard deviation. Statistical significance of changes in MR parameters is marked with asterisks (ns: *p* > 0.05, *: *p* < 0.05, **: *p* < 0.01, ***: *p* < 0.001).

The effects of Δ*V*_*m*_ on *T*_2_ and PSR were both significant (*p* < 0.0001), indicating an increase in *T*_2_ and a decrease in PSR during depolarization at high [K^+^], with the opposite trend during hyperpolarization at low [K^+^]. On the other hand, the effect of Δ*V*_*m*_ on *k*_*mf*_ was not significant (*p* = 0.360).

Subsequent post-hoc analyses compared each experimental condition to the control using Dunnett’s test to account for multiple comparisons (Fig. 2). During depolarization induced by the highest [K^+^] (80 mM, Δ*V*_*m*_ = 30.0 mV), changes in MR parameters were observed as a 20.0% increase in *T*_2_ (Δ*T*_2_ = 10.1 ms, *p* < 0.0001) and a 12.9% decrease in PSR (ΔPSR = −0.00476, *p* < 0.0001). Conversely, during hyperpolarization induced by the lowest [K^+^] (0.2 mM, Δ*V*_*m*_ = −5.33 mV), *T*_2_ decreased by 4.40% (Δ*T*_2_ = −2.21 ms, *p* < 0.0001) and PSR increased by 6.28% (ΔPSR = 0.00231, *p* < 0.0005). However, changes in *k*_*mf*_ were not significant across all conditions (*p* > 0.05). These findings from *in vitro* SH-SY5Y cell experiments suggest that MR parameters, such as *T*_2_ and PSR, exhibit sufficient sensitivity to detect alterations in membrane potential induced by varying [K^+^], including both depolarization and hyperpolarization.

### Using a K^+^ channel blocker

In this experiment, we investigated whether depolarization induced by altering potassium permeability with barium ions (Ba^2+^) would affect MR parameters similarly to depolarization induced by varying [K^+^], thereby further validating our findings. For this purpose, we administered barium ions (Ba^2+^) at a concentration of 10 mM to induce depolarization while maintaining constant osmolarity by adjusting [Na^+^]. Ba^2+^ was chosen because it inhibits several types of two-pore-domain potassium channels^37,38^, which predominantly regulate the resting membrane potential. Previous studies have confirmed that Ba^2+^ depolarizes the membrane potential in SH-SY5Y and Jurkat cells^39,40^.

The Ba^2+^-induced depolarization condition was compared with K^+^-induced depolarization and hyperpolarization conditions (Fig. 3). To compare the effects of [Ba^2+^] and [K^+^] on MR parameters, a linear mixed-effect model was utilized. This model included MR parameters as dependent variables, Δ*V*_*m*_ and its interaction with a group variable (indicating whether Δ*V*_*m*_ was induced by [Ba^2+^] or [K^+^]) as fixed effects, and cell batch as a random effect. This analysis revealed no significant interactions for all MR parameters assessed (*p* = 0.182 for *T*_2_, *p* = 0.788 for PSR, and *p* = 0.0890 for *k*_*mf*_). These findings suggest that changes in *T*_2_ and PSR by membrane potential do not depend on the specific method of altering the membrane potential, whether by varying [K^+^] or applying [Ba^2+^]. This implies that if the changes in MR parameters observed with varying [K^+^] were not primarily due to membrane potential but were due to unique K^+^-related mechanism, then experiments using [Ba^2+^] would not result in similar changes.

**Fig. 3.**
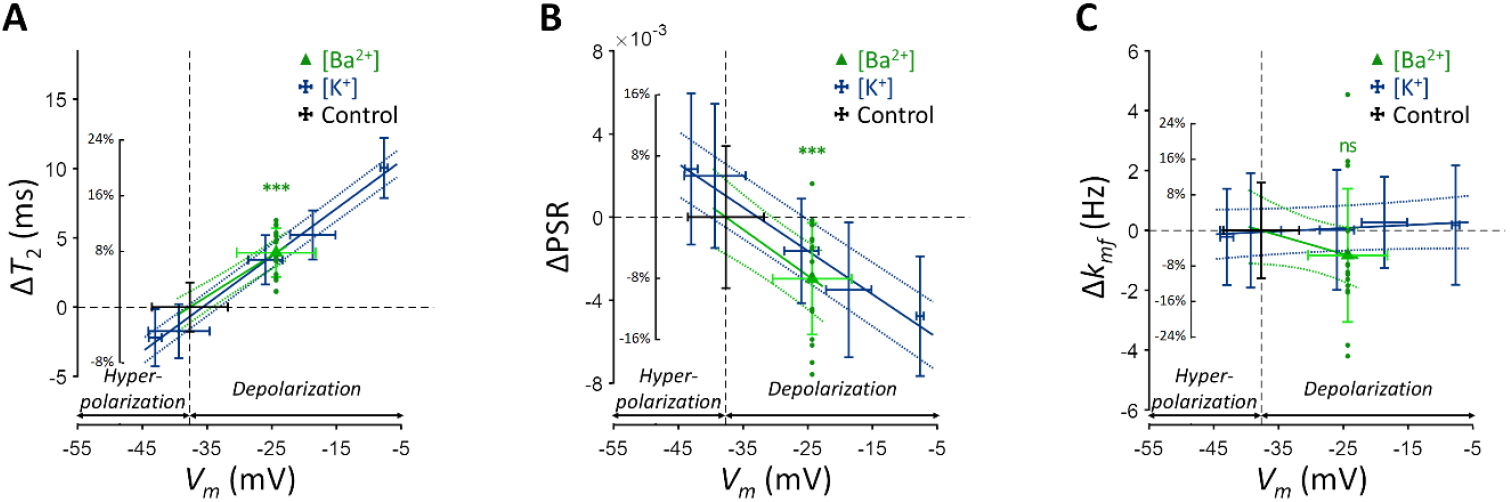
Changes in (A) *T*_2_, (B) PSR, and (C) *k*_*mf*_ of SH-SY5Y cells across experimental conditions: [K^+^] = 0.2–80 mM (blue cross) and [Ba^2+^] = 10 mM (green triangle), compared to the control condition (black cross). Data from fifteen experiments (*n* = 15) are displayed. Linear regression lines for [K^+^] data (blue solid line) and [Ba^2+^] data (green solid line) are drawn along with dotted lines representing 95% confidence intervals. Error bars denote standard deviation. Statistical significance of changes in MR parameters with [Ba^2+^] = 10 mM is marked with asterisks (ns: *p* > 0.05, *: *p* < 0.05, **: *p* < 0.01, ***: *p* < 0.001).

Subsequent post-hoc analyses compared the [Ba^2+^]-induced depolarization to the control using Dunnett’s test to account for multiple comparisons (Fig. 3). In response to depolarization caused by [Ba^2+^] = 10 mM, *T*_2_ increased by 7.82% (Δ*T*_2_ = 3.93 ms, *p* < 0.0001) and PSR decreased by 8.06% (ΔPSR = −0.00297, *p* < 0.0001). The change in *k*_*mf*_ was not significant (Δ*k*_*mf*_ = −0.832 Hz, *p* = 0.263). The depolarization of membrane potential induced by [Ba^2+^] was measured as Δ*V*_*m*_ = 13.3 mV by patch clamp recording.

### Using another cell type

To investigate whether membrane potential-modulating ionic solutions produce similar changes in MR parameters across different cell types, we assessed another cell line, Jurkat, under the same experimental conditions applied to SH-SY5Y cells. These conditions included a control condition ([K^+^] = 4.2 mM), hyperpolarization under decreased [K^+^] conditions ([K^+^] = 0.2 and 1 mM), depolarization under increased [K^+^] conditions ([K^+^] = 20, 40, and 80 mM), and a Ba^2+^-induced depolarization condition ([Ba^2+^] = 10 mM), all maintaining consistent osmolarity by adjusting [Na^+^].

Each experimental condition was compared to the control using Dunnett’s test to account for multiple comparisons (Fig. 4). As observed with SH-SY5Y cells, Jurkat cells showed significant positive changes in *T*_2_ and negative changes in PSR under increased [K^+^] conditions. For example, at [K^+^] = 80 mM, *T*_2_ increased by 16.9% (Δ*T*_2_ = 9.26 ms, *p* < 0.0001), PSR decreased by 21.9% (ΔPSR = −0.00347, *p* < 0.01). In contrast, during hyperpolarization at the lowest [K^+^] = 0.2 mM, *T*_2_ decreased by 18.1% (Δ*T*_2_ = −9.93 ms, *p* < 0.0001), whereas PSR increased by 17.6% (ΔPSR = 0.00280, *p* < 0.05). The depolarization induced by [Ba^2+^] = 10 mM resulted in a similar pattern, with *T*_2_ increasing by 11.5% (Δ*T*_2_ = 6.3 ms, *p* < 0.0005), although the decrease in PSR was not significant (ΔPSR = −0.00212, *p* = 0.211). Changes in *k*_*mf*_ remained non-significant across all conditions (*p* > 0.05). In summary, these findings indicate that detecting membrane potential changes induced by ionic solutions using MR parameters such as *T*_2_ and PSR is not specific to a single cell type, although the magnitude of these changes may differ between cell types.

**Fig. 4.**
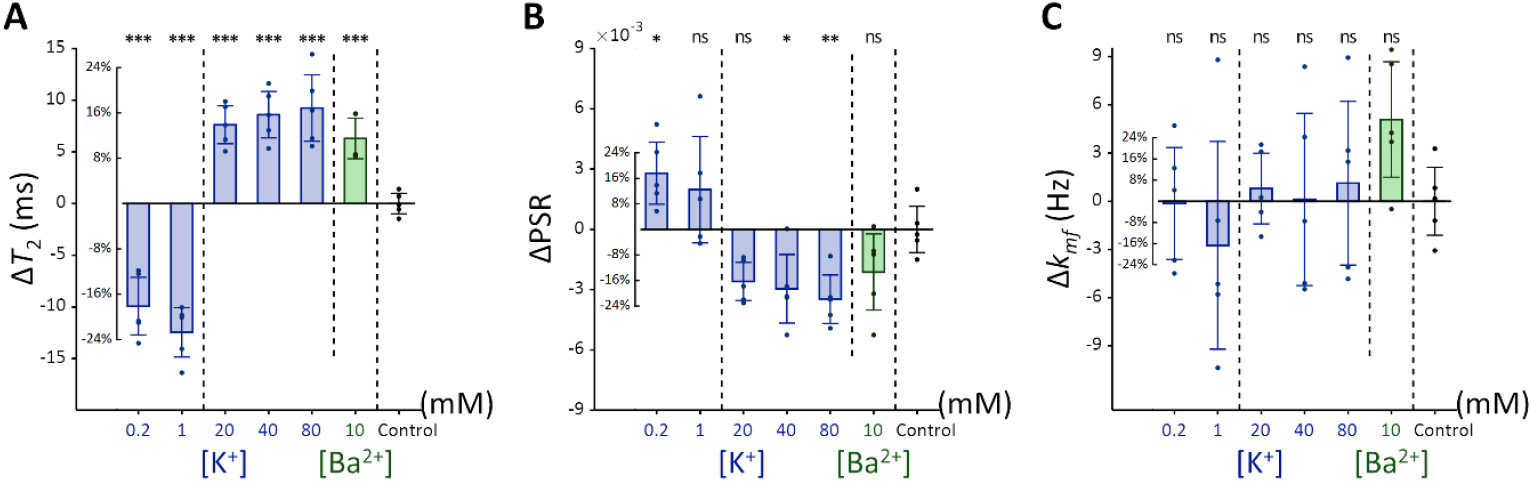
Changes in (A) *T*_2_, (B) PSR, and (C) *k*_*mf*_ of Jurkat cells across experimental conditions: [K^+^] = 0.2–80 mM (blue bar) and [Ba^2+^] = 10 mM (green bar), compared to the control condition of [K^+^] = 4.2 mM (*n* = 5). Error bars denote standard deviation. Statistical significance of changes in MR parameters is marked with asterisks (ns: *p* > 0.05, *: *p* < 0.05, **: *p* < 0.01, ***: *p* < 0.001).

### *In vivo* changes in *T*_2_ by membrane potential

The relationship of *T*_2_ values and membrane potential modulated by [K^+^], observed in the aforementioned SH-SY5Y and Jurkat cell studies, was further explored in an *in vivo* rat model to validate these findings under physiological conditions. As depicted in Figure 5, a craniotomy was performed to expose a 3-mm-diameter region of the rat cerebral cortex, followed by perfusion with artificial cerebrospinal fluid (aCSF) to modulate the membrane potential. Hemodynamic responses were pharmacologically suppressed. MRI scans were performed using a 7 T preclinical MRI system to measure *T*_2_ in the exposed cortical area. A total of seven rats were used in the experiment that involved modulation of [K^+^]. The experimental protocol included sequential application of four conditions, each lasting 12 minutes: a baseline condition at [K^+^] = 3 mM, a depolarization condition at [K^+^] = 40 mM, further depolarization at [K^+^] = 80 mM, followed by a recovery condition using baseline aCSF. The recovery condition was applied to two of the seven rats. To distinguish the effect of aCSF perfusion on *T*_2_ from the effect of changes in membrane potential, a control experiment was also conducted using only baseline aCSF for the entire duration (48 minutes) with another group of five rats.

**Fig. 5.**
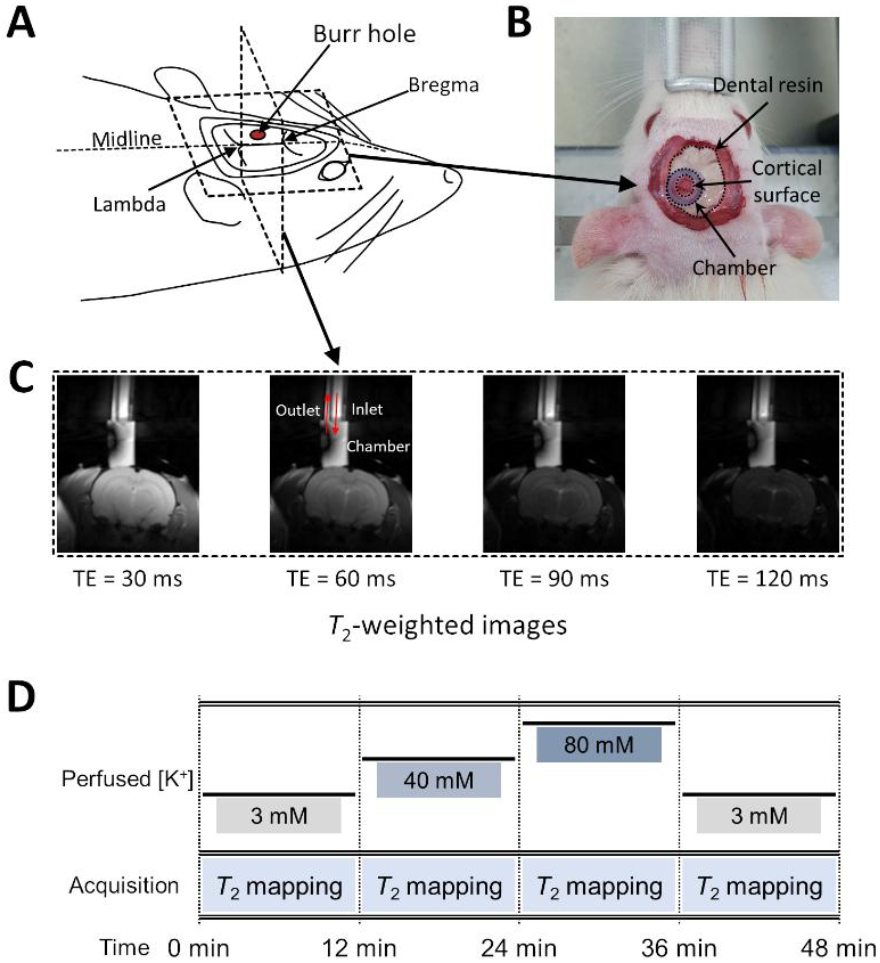
Experimental setup for *in vivo* manipulation of membrane potential. (A) A schematic diagram of the rat head post-craniotomy, showing the burr hole centered at 2.5 mm anterior and 2.0 mm lateral to the lambda. (B) Photograph of the rat head with a cylindrical chamber fixed over the burr hole, filled with artificial cerebrospinal fluid. (C) A representative series of *T*_2_-weighted MR images for *T*_2_ mapping. The chamber was connected to inlet and outlet perfusion tubes. (D) The experimental paradigm of the *in vivo* rat MR imaging. Four conditions were sequentially applied: control, depolarization, further depolarization, and recovery. Each condition lasted 12 minutes during which *T*_2_ mapping were conducted.

In Figure 6A, a representative *T*_2_ map is displayed with an enlarged image defining the ROI beneath the perfusion chamber (width = 1.8 mm, depth = 0.6mm). The average *T*_2_ value within this ROI was estimated from the spatially averaged multi-echo spin-echo signal. A detailed analysis on the quality of these *T*_2_ maps is presented in Supplementary Results S2.1. Changes in *T*_2_ value (Δ*T*_2_) were statistically analyzed using a linear mixed-effect model to account for inter-sample variability and repeated measurements. This model included Δ*T*_2_ as a dependent variable, elapsed time and its interaction with the experiment type (i.e., [K^+^]-modulation or control) as fixed effects, and a random effect for inter-sample variability.

**Fig. 6.**
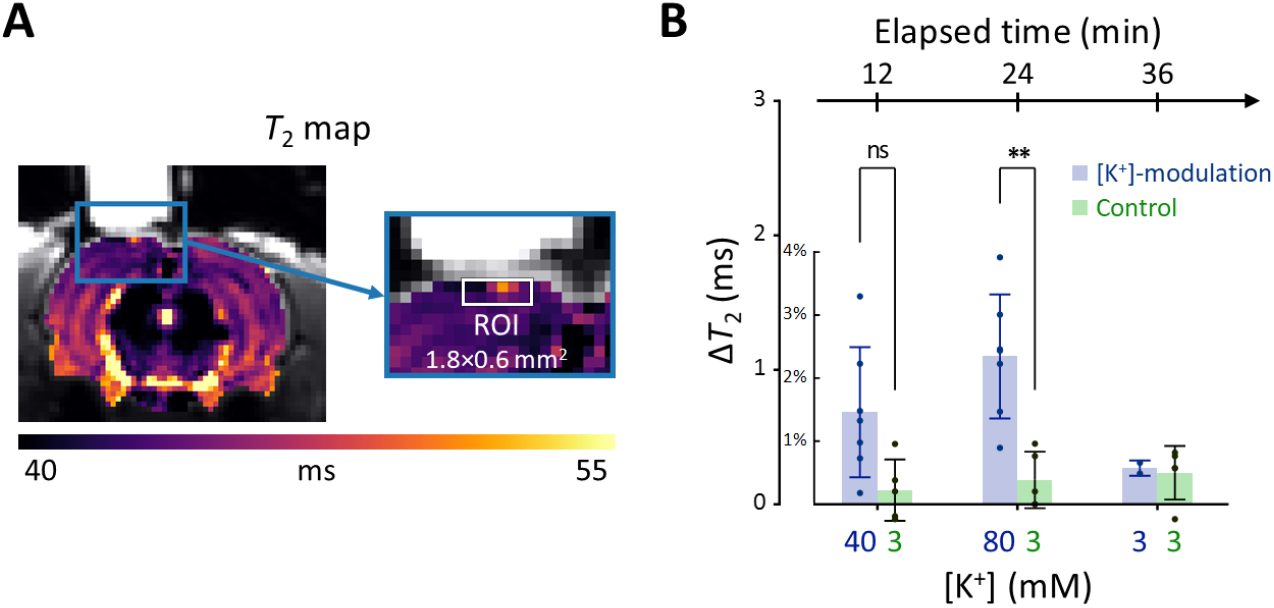
Results of the *in vivo* experiment results in rat models. (A) A representative *T*_2_ map from a single rat with an enlarged image depicting the ROI for estimating average *T*_2_ in the exposed cortical area, marked by a white rectangle (width = 1.8 mm, depth = 0.6 mm). (B) The changes in *T*_2_ values within the ROI is plotted against elapsed time from the initial conditions. [K^+^] in the perfused artificial cerebrospinal fluid are indicated on the bottom abscissa. Results from the [K^+^]-modulation experiments are shown with blue bars, and those from the control experiments are shown with green bars. Statistical significance of the *T*_2_ changes are marked with asterisks (ns: *p* > 0.05, *: *p* < 0.05, **: *p* < 0.01).

Figure 6B shows the results of the statistical analysis. Δ*T*_2_ values were plotted against elapsed time for [K^+^]-modulation and control experiments. After 12 minutes from the initial condition, *T*_2_ increased by 1.46% (Δ*T*_2_ = 0.684 ms) in the [K^+^]-modulation experiment ([K^+^] = 40 mM) and by 0.223% (Δ*T*_2_ = 0.104 ms) in the control experiment ([K^+^] = 3 mM); the Δ*T*_2_ difference (= 0.580 ms) between these experiments was not statistically significant (*p* = 0.0711). After 24 minutes, *T*_2_ increased by 2.34% (Δ*T*_2_ = 1.10 ms) in the [K^+^]-modulation experiment ([K^+^] = 80 mM) and by 0.386% (Δ*T*_2_ = 0.181 ms) in the control experiment ([K^+^] = 3 mM), with a significant difference in Δ*T*_2_ (= 0.918 ms) between them (*p* = 0.00172). Due to the limited sample size in the recovery phase (*n* = 2 out of 7 for [K^+^]-modulation), comparisons between experiments were not conducted for this recovery phase.

Our observations indicate that *in vivo* manipulation of membrane potential results in similar trends of *T*_2_ changes as those observed *in vitro*, albeit with a smaller magnitude in the rat cortex. This discrepancy may be attributed to several factors: [K^+^]-modulation affecting only a small portion of cells within the ROI; limited diffusion of aCSF through the leptomeninges; removal of excessive K^+^ through clearing mechanisms; and differences in cell types (see the Discussion for more details).

## Discussion

In this study, we demonstrated that MR parameters, specifically *T*_2_ relaxation time and pool size ratio (PSR), can detect responses to membrane potential changes modulated by ionic solutions. Our *in vitro* experiments with cultured cells were designed to exclude physiological factors such as hemodynamic responses and respiration. We observed that depolarization increases *T*_2_ and decreases PSR, while hyperpolarization has the opposite effect. *In vivo*, we pharmacologically suppressed hemodynamic responses to minimize their impact on *T*_2_ measurements. The trend of *T*_2_ dependence on membrane potential *in vivo* was consistent with *in vitro* findings. However, the magnitude of *T*_2_ changes in rat cortex was approximately one-ninth of that observed in SH-SY5Y cells *in vitro* at [K^+^] = 80 mM.

Other studies have also reported MR-detectable changes in response to extracellular [K^+^] modulation. For instance, research on spreading depression, which was induced by significantly high [K^+^] (∼1 M) applied topically to rat cortex, has revealed detectable changes in *T*_1_, *T*_2_, and magnetization transfer ratio changes^41^; and spin-lock fMRI signals^42^. Similarly, increased [K^+^] has been studied *in vitro* with brain slices, linking cell volume changes to proton density-weighted MR signal changes^43^. *T*_2_ mapping in Jurkat cells with increased [K+] conditions has also been investigated^27^ and linked to cell volume change^44^. Our study complements these findings by employing direct membrane potential measurement via patch clamp, testing additional ionic agents such as Ba^2+^, and demonstrating the phenomenon *in vivo*. Further reinforcement of these findings could be achieved by simultaneous recording of cell volume and other cellular characteristics to elucidate the underlying mechanisms more completely.

Interestingly, the MR responses in Jurkat cells differed from those in SH-SY5Y cells. While SH-SY5Y cells showed a near-linear dependence of MR parameters on log [K^+^], Jurkat cells displayed more step-like behavior. The reasons for this discrepancy remain unclear, but it suggests that the relationship between membrane potential and MR parameters may not strictly follow a simple log [K+] relationship.

Several factors may contribute to discrepancies between *in vivo* and *in vitro* results. For instance, the actual extracellular [K^+^] experienced by cells in the rat cortex may be lower than that of the perfused aCSF due to diffusion-limiting barriers such as the leptomeninges^45,46^, even after the removal of the dura mater. Additionally, removal of excessive K^+^ by clearing mechanisms^47,48^ may further reduce the [K^+^] experienced by the cells. Partial volume effects and cell type differences may also contribute to the discrepancy.

From a biophysical perspective, the sensitivity of *T*_2_ and PSR to membrane potential likely arises from alterations in cell volume, hydration water, and bulk water. Depolarization or hyperpolarization can lead to cell swelling or shrinking^49-51^, influencing the proportion of cellular contents within the imaging voxel and thereby affecting MR parameters. Notably, neuronal or glial cell swelling has been proposed as a possible mechanism underlying diffusion fMRI^29,52,53^ and previous MRI studies^43,44^. Although not as sensitive as *T*_2_, our supplementary *T*_1_ measurements (Supplementary Fig. S2–4) also exhibited similar trends in response to membrane potential changes. Moreover, hydration water, which has a significantly shorter *T*_2_ than bulk water due to slower re-orientational and diffusive motions^54^, may contribute to the observed MR changes. In particular, the correlation between PSR and membrane potential indicates that depolarization decreases the density of hydration water on the cell membrane within a voxel, thereby reducing PSR and simultaneously increasing *T*_2_ due to a corresponding increase in free water, as well as cell swelling. Conversely, hyperpolarization may increase hydration water density, elevating PSR and lowering *T*_2_. These interpretations are supported by recent optical studies showing reduced membrane hydration water during depolarization^55^.

Several challenges and considerations in this study warrants discussion. First, maintaining the desired environment (37°C and 5% CO_2_) for *in vitro* cells during MRI scans is challenging. In addition, intracellular accumulation of Ba^2+^ in the Ba^2+^-induced depolarization experiment may affect cellular integrity. In this study, high cell viability (> 97.6%) was confirmed using cell viability assays under experimental conditions (Supplementary Fig. S5). Second, differences in *T*_2_ value among the extracellular media may bias *T*_2_ measurements due to the partial volume effect. However, in this study, the differences in *T*_2_ among the extracellular media were found to be negligible compared to the observed *T*_2_ changes (Supplementary Fig. S6). Third, while our findings show that membrane potential-modulating ionic solutions can affect MR parameters, it is important to note that these changes do not measure the membrane potential itself. Fourth, other factors such as pH, energy depletion, or extracellular osmolarity may affect MR parameters by altering cell volume. To minimize the effects of these other contributors, we matched osmolarity across all conditions, provided sufficient glucose to prevent energy depletion, and regulated pH levels by buffering with HEPES. Additionally, to mitigate possible changes in intra/extracellular volume fraction changes caused by cell movements, potentially due to agitation, we centrifuged the cells and imaged the bottom portion of the cell pellet, where cell movement was restricted due to close packing. Fifth, in the *in vivo* study, we attempted to suppress hemodynamic responses through pharmacological means, using a combination of N_ω_-Nitro-L-arginine and nifedipine, both of which are known to inhibit hemodynamic responses in different ways^56,57^, but their effects were not directly confirmed in this study. Future studies that simultaneously evaluate hemodynamic responses would strengthen our conclusions. Finally, our experimental paradigm was based on clamping the membrane potential at a specific level, thus measuring changes in *T*_2_ and PSR during static depolarization or hyperpolarization rather than dynamic changes such as those seen during action potentials. Future research could explore temporally varying membrane potential to evaluate the dynamic correlation between membrane potential and MR parameters with high temporal resolution MRI^26,27^.

In summary, our study demonstrates that MR parameters such as *T*_2_ relaxation time can detect responses to membrane potential-modulating ionic solutions both *in vitro* and *in vivo*. This finding proposes a potential approach for noninvasively detecting changes in membrane potential using MRI.

## Methods

### *In vitro* cell culture

Two human cell lines were utilized for the experiments: SH-SY5Y, an immortalized neuroblastoma line, and Jurkat, a leukemia cell line. Both cell lines were sourced from the American Type Culture Collection (ATCC). The SH-SY5Y cells were cultured in DMEM/F12 medium supplemented with 10% (v/v) fetal bovine serum (FBS) and 100 U/ml penicillin/streptomycin. The cells were maintained at a constant temperature of 37°C in a humidified atmosphere containing 5% CO_2_. Similarly, the Jurkat cells were cultured in RPMI-1640 medium, also supplemented with 10% (v/v) FBS and 100 U/ml penicillin/streptomycin, under the same conditions of temperature and CO_2_ concentration.

### *In vitro* manipulation of membrane potential with extracellular media

The baseline extracellular medium was prepared with the following components: KCl = 4.2 mM; NaCl = 145.8 mM; HEPES = 20 mM; glucose = 4.5 g/l; EGTA = 10 µM; pH = 7.2. This baseline medium was considered a control condition for various extracellular media used to adjust membrane potential. Two extracellular media with low K^+^ concentrations (KCl = 0.2 and 1 mM) were prepared to hyperpolarize the membrane potential. Three extracellular media with high K^+^ concentrations (KCl = 20, 40, and 80 mM) were prepared to depolarize the membrane potential. A Ba^2+^ medium containing 10 mM BaCl_2_ was also prepared to depolarize the membrane potential in a different way, i.e., as a K^+^ channel blocker. These seven extracellular media were used for both MR imaging and patch clamp recording *in vitro*. Sodium ion concentrations ([Na^+^]) in all media were controlled to match the osmolarity with the baseline medium. The composition of all extracellular media is detailed in Supplementary Table S3.

### Preparation of cells for *in vitro* MR measurement

SH-SY5Y cells were dissociated from their culture plates using 0.5 mM EDTA solution, then concentrated into a pellet (∼70 µl) by centrifugation at 250 × g for two minutes. The pellet was resuspended in the culture medium and divided evenly into seven aliquots. Each cell suspension was centrifuged and resuspended in each of the seven different media specified in Supplementary Table S3. Centrifugation and resuspension were repeated three more times to completely clear out the culture medium. Each cell suspension with a different extracellular medium was then loaded into two wells on the same column of an acrylic container with 14 wells (matrix = 2 × 7) and centrifuged again to concentrate into pellets (Fig. 1). The acrylic container was purposely formed into a spherical segment to enhance the homogeneity of the static magnetic field^58^. The preparation of Jurkat cell samples was the same as for the SH-SY5Y cell samples. The preparation required 40–60 minutes, followed by an incubation period of 20–30 minutes.

### *In vitro* MRI experiment

*In vitro* MRI experiments were performed on a 9.4 T MRI system (BioSpec 94/30 USR, Bruker BioSpin) at room temperature. A volume coil with an inner diameter of 86 mm was utilized for both radiofrequency (RF) pulse transmission and signal reception. Within the acrylic container, two wells on the same column (matrix = 2 × 7) contained identical cell pellets with the same extracellular media. MRI signals from seven different cell samples in the horizontal direction were separated by one-dimensional frequency encoding along that direction (Fig. 1). The MRI pulse sequence employed for mapping the *T*_2_ value was a single-echo spin-echo (SESE) sequence with 50 variable echo times (TE) spaced between 9.5 and 290.5 ms on a logarithmic scale. For mapping the MT parameters, an inversion recovery multi-echo spin-echo (IR-MESE) sequence was used. The inversion times (TI) for the IR-MESE sequence were optimized using the theory of Cramér-Rao lower bounds^59^. The optimized TIs ranged from 4 to 10,079.4 ms. After each TI, 16 spin-echo trains were acquired with an echo spacing of 9.5 ms. Total scan time for both sequences was 23 minutes. Experiments were repeated 15 times for SH-SY5Y cells and 7 times for Jurkat cells, replacing cells in each repetition. Other scan parameters are detailed in Supplementary Table S1.

### Animals

Male Wistar rats aged 8 weeks (250–300 g, Orient Bio) were used for MRI experiments after undergoing a craniotomy. All animal experiments were approved by the Institutional Animal Care and Use Committee at the National Cancer Center Korea (NCC-22-740). The rats were housed in ventilated cages under a 12h/12h light/dark cycle and provided with *ad libitum* access to food and water.

### *In vivo* manipulation of membrane potential with artificial cerebrospinal fluid (aCSF)

The membrane potential of the exposed cortex was manipulated by directly perfusing the region-of-interest of the cerebral cortex with aCSF after a craniotomy. The baseline aCSF was prepared with the following components: KCl = 3 mM; NaCl = 135 mM; MgCl_2_ = 3 mM; HEPES = 20 mM; glucose = 4.5 g/l; EGTA = 2 mM; N_ω_-Nitro-L-arginine = 1 mM; Nifedipine = 0.1 mM; pH = 7.4. Hemodynamic effects were pharmacologically suppressed using N_ω_-Nitro-L-arginine, Nifedipine, and EGTA. N_ω_-Nitro-L-arginine suppresses depolarization-induced hemodynamic response by blocking the synthesis of nitric oxide, which acts as a vasodilator^56^. Nifedipine blocks voltage-sensitive Ca^2+^ channels, and EGTA chelates free Ca^2+^ to inhibit the hemodynamic response^57^. To induce depolarization, aCSF with high [K^+^] (KCl = 40 and 80 mM) was prepared. The osmolarity of the aCSF was matched with the concentration of NaCl.

### Rat surgery

A craniotomy was performed on a Wistar rat. The experimental setup after a surgical procedure is illustrated in Figure 5. The surgery was performed following an established protocol^60^ alike to that used in other MRI studies^41,42^. Anesthesia was induced with 3% isoflurane in O_2_ and maintained with 2–3% isoflurane during the surgical procedure. Body temperature was maintained at 36.5–37.5°C with an infrared lamp. The head was fixed with a small animal stereotaxic frame. The hair on the scalp was shaved with a veterinary clipper. The skin and periosteum over the skull were removed using a scalpel and surgical scissors. A 3.0-mm-diameter burr hole was opened using a dental drill with its center at the coordinates of 2.5 mm anterior and 2.0 mm lateral to the lambda. Then, a cylindrical chamber was implanted upon the burr hole with cyanoacrylate glue and dental composite resin. The chamber was filled with the baseline aCSF and connected to inlet and outlet perfusion tubes. The exposed cerebral cortex inside the chamber was perfused with the baseline aCSF at a flow rate of 0.6 ml/min using peristaltic pumps.

### *In vivo* MRI experiment

The rat with a cranial chamber installed on the cortical surface was placed on a 7 T MRI system (BioSpec 70/20 USR, Bruker BioSpin), fixed in a customized cradle with two ear-bars and a bite-bar. A customized surface coil (rectangular, 35 mm × 20 mm) was used for RF pulse transmission and signal reception. Body temperature was maintained at 36.5–37.5°C using a warm air blower. Anesthesia was maintained with 2% isoflurane in O_2_ (0.6 l/min). Respiration rate and body temperature were monitored throughout the MRI experiment. MR images were acquired in a 2 mm coronal slice through the center of the burr hole (Fig. 5), using a multi-echo spin-echo (MESE) sequence with 20 TEs (7.5 to 150 ms). Other scan parameters are detailed in Supplementary Table S2.

A total of seven rats were subjected to four sequential experimental conditions, as depicted in Figure 5. First, the exposed cerebral cortex was perfused with baseline aCSF ([K^+^] = 3 mM). Second, as a depolarizing condition, the perfusion media was switched to depolarizing aCSF of [K^+^] = 40 mM. Third, membrane potential was further depolarized by perfusion with aCSF of [K^+^] = 80 mM. Finally, as a recovery condition, the perfused aCSF was changed back to baseline aCSF. During each condition, MR images were acquired with MESE sequences for 12 minutes. Two rats underwent the whole four conditions, while other five rats did not undergo the recovery condition. As a control experiment, another set of rats (*n* = 5) underwent perfusion with the baseline aCSF for the same duration (48 minutes) as the previous experiment, and MR images were acquired with four MESE sequences, each for 12 minutes. Throughout the experiment, the perfusion rate was maintained at 0.6 ml/min.

### Quantification of MR parameters

For the *in vivo* MR images acquired with a MESE sequence, echo trains were matched with a simulated dictionary of decay curves of multi-echo spin-echo signals created with the stimulated echo and slice profile correction^61^ to estimate *T*_2_ values. For the *in vitro* one-dimensional MR images acquired with the SESE sequence, signals were fitted to a mono-exponential function to estimate *T*_2_ values. For the *in vitro* one-dimensional MR images acquired with the IR-MESE sequence, signals were fitted to a bi-exponential function^62,63^ to estimate MT parameters. 16 spin echoes acquired after each TI were averaged to improve SNR. The MT parameters such as PSR and *k*_*mf*_ were derived from this fitting process. The detailed procedure for calculating *T*_2_ and two MT parameters (i.e., PSR and *k*_*mf*_) is provided in Supplementary Methods S1.1.

### Patch clamp recordings

The membrane potential of SH-SY5Y cells was recorded at room temperature using the whole-cell mode of the patch clamp technique^64^. The bath solution was the same as the extracellular medium used in the MRI experiment and was constantly perfused at a flow rate of 2 ml/min. The composition of the pipette solution was as follows: KCl = 140 mM; NaCl = 5 mM; MgCl2 = 3 mM; HEPES = 10 mM; Mg-ATP = 1 mM; Na-GTP = 0.5 mM. The pH of the pipette solution was adjusted to 7.4 using KOH. Calcium ions (Ca^2+^) were not included in the pipette solution to minimize Ca^2+^-dependent currents. The resistance of the electrode was 3–5 MΩ with the internal solution filled. Recordings were performed with a patch amplifier (Axopatch-1D; Axon Instruments) and a current clamp was also used to monitor the membrane potential. The experiment was repeated three times, with cells replaced at each repetition.

## Acknowledgements

We thank Dr. Silvia Mangia for insightful discussions on the role of hydration water in functional MRI. Animal molecular imaging facility at the National Cancer Center Korea contributed for supportive animal magnetic resonance imaging. We thank Minsun Kim and Soyeon Jeon (National Cancer Center) for helping with rat craniotomies and MRI experiments. This work was supported by the National Research Foundation of Korea grant funded by the Ministry of Science and ICT (NRF-2019M3C7A1031993, NRF-2019M3C7A1031994, and NRF-2023R1A2C3007075). This work was supported by the Commercialization Promotion Agency for R&D Outcomes (COMPA) funded by the Ministry of Science and ICT (NTIS1711198890).

## Author contributions

**Conceptualization**: K.M., S.-K.L., and J.-Y.P. **Methodology**: K.M., S.C., S.-K.L., J.L., P.T.T., D.K., J.S.L., and J.-Y.P. **Investigation**: K.M., S.C., P.T.T., and D.K.; **Resources**: S.C., J.L., D.K., J.S.L., and J.-Y.P. **Visualization**: K.M., and S.C. **Supervision**: J.L., and J.-Y.P. **Writing – original draft** K.M., and J.-Y.P. **Writing – review & editing**: K.M., S.C., S.-K.L., J.L., P.T.T., D.K., J.S.L., and J.- Y.P.

## Data availability

The data that support the findings of this study are available from the corresponding author upon reasonable request.

