## Supplementary Information for "Responses to membrane potential-modulating ionic solutions measured by magnetic resonance imaging of cultured cells and *in vivo* rat cortex"

### Table of contents

**Supplementary Methods**
S1.1. Quantification of $T_1$ and MT parameters
S1.2. Quantification of $T_1$ and $T_2$ of extracellular media
S1.3. Viability assay of SH-SY5Y cells with live/dead staining

**Supplementary Results**
S2.1. Quantitative assessment of *in vivo* $T_2$ mapping
S2.2. Membrane potential-induced changes in $T_1$: SH-SY5Y cells
S2.3. Membrane potential-induced changes in $T_1$: Jurkat cells

**Supplementary Figures**
• Figures S1–S6

**Supplementary Tables**
• Tables S1–S8

**Supplementary References**

### Supplementary Methods

#### S1.1. Quantification of $T_1$ and MT parameters

To estimate the $T_1$ and MT parameters, MR signals acquired using IR-MESE sequences were fitted to a bi-exponential function[1,2]. 16 spin echoes obtained after each TI were averaged to improve the SNR. The MT parameters derived from this fitting process include the pool size ratio (PSR) and the magnetization transfer rate from macromolecules to free water ($k_{mf}$). A 1 ms hard inversion pulse selectively inverted the magnetization of free water protons, leading to cross-relaxation between the longitudinal magnetization of free water protons ($M_{z,f}$) and macromolecular protons ($M_{z,m}$). This interaction resulted in a bi-exponential magnetization recovery characterized by a fast longitudinal relaxation rate $R_1^+$ and a slow relaxation rate $R_1^-$ (= $1/T_1$), with the latter corresponding to the conventional spin-lattice relaxation rate[2,3]:

$$\frac{M_{z,f}(t)}{M_{\infty,f}} = b_f^+ \exp(-R_1^+ t) + b_f^- \exp(-R_1^- t) + 1 \qquad [S1]$$

where $M_{\infty,f}$ denotes the equilibrium magnetization of free water protons, and $b_f^+$ and $b_f^-$ denote the amplitudes for the exponential terms associated with $R_1^+$ and $R_1^-$, respectively. By fitting this bi-exponential model to the inversion recovery signals using least squares, estimates of $R_1^+$, $R_1^-$, $b_f^+$, and $b_f^-$ were obtained. These parameters are related to the MT parameters, PSR and $k_{mf}$, by following equations:

$$2R_1^{\pm} = R_{1,f} + R_{1,m} + k_{fm} + k_{mf} \pm \sqrt{\left(R_{1,f} - R_{1,m} + k_{fm} - k_{mf}\right)^2 + 4k_{fm}k_{mf}} \qquad [S2]$$

$$b_f^{\pm} = \pm \frac{\left(R_{1,f} - R_1^{\mp}\right)\left(\frac{M_{z,f}(0)}{M_{\infty,f}} - 1\right) + k_{fm}\left(\frac{M_{z,f}(0)}{M_{\infty,f}} - \frac{M_{z,m}(0)}{M_{\infty,m}}\right)}{R_1^+ - R_1^-} \qquad [S3]$$

$$PSR = k_{fm}/k_{mf} \qquad [S4]$$

where $R_{1,f}$ and $R_{1,m}$ denote the longitudinal relaxation rates of free water and macromolecular protons, respectively, in the absence of cross-relaxation. $k_{fm}$ and $k_{mf}$ denote the magnetization transfer rates from free water to macromolecules and vice versa, respectively, $M_{z,m}$ (0) denote the longitudinal magnetization of macromolecular protons immediately after the inversion pulse. $M_{\infty,m}$ denote the equilibrium magnetization of macromolecular protons. According to previous studies[2,4], $M_{z,m}(0)/M_{\infty,m}$ can be determined numerically by the Bloch equations. Assuming $R_{1,f} = R_{1,m}$[5,6], simplifies the equations [S2–4] to explicitly calculate PSR and $k_{mf}$:

$$PSR = \frac{b_f^+}{b_f^- - \frac{M_{z,m}(0)}{M_{\infty,m}} + 1} \qquad [S5]$$

$$k_{mf} = \frac{R_1^+ - R_1^-}{1 + PSR} \qquad [S6]$$

### S1.2. Quantification of $T_1$ and $T_2$ of extracellular media

To estimate the $T_1$ and $T_2$ of extracellular media specified in Supplementary Table S3, 200 µl of each media sample was loaded into two wells on the same column of the acrylic container with 14 wells (Fig. 1A) and placed in a 9.4 T preclinical MRI system (BioSpec 94/30 USR, Bruker BioSpin). $T_1$ and $T_2$ were measured using an adiabatic inversion recovery multi-echo spin-echo sequence. 20 variable inversion times (TI) were spaced between 600 and 15,000 ms on a logarithmic scale. After each TI, 50 multi-echo spin-echo trains were acquired with an echo spacing of 9.5 ms (echo time (TE) = 9.5, 19, …, 475 ms). For the $T_1$ estimation, the 50 multi-echo spin-echo signals acquired after each TI were averaged and fitted to a mono-exponential function:

$$S(t) = A + B\exp(-t/T_1) \quad \text{[S7]}$$

where $S(t)$ denotes the averaged signal after an inversion time of $t$, and A and B denote the coefficients of the fitting. For $T_2$ estimation, the 20 spin-echo signals at the same TE but different 20 TIs were averaged. These averaged 50 multi-echo spin-echo trains were then matched with a simulated dictionary of decay curves. This dictionary was constructed from simulation of multi-echo spin-echo signals, incorporating corrections for stimulated echoes and slice profile effects[7].

### S1.3. Viability assay of SH-SY5Y cells with live/dead staining

SH-SY5Y cells were cultured, harvested and centrifuged to obtain a cell pellet. This cell pellet was resuspended in the culture medium and divided into seven equal aliquots. Each aliquot was centrifuged and resuspended three times using one of the seven different media specified in Supplementary Table S3. For the viability assay, staining media were prepared by adding 2 µM calcein-AM and 4 µM Ethidium homodimer-1 to each of the seven media. The cell suspensions were then centrifuged and resuspended again using the staining media. Following this, each cell suspension was transferred to separate wells in a chambered coverglass and incubated at room temperature for 20 minutes. After another round of centrifugation, cell pellets were obtained and analyzed using a confocal microscope (Leica TCS SP8 STED, Leica Microsystems). Cell viability was assessed based on green fluorescence indicating live cells and red fluorescence indicating or dead cells, under excitation wavelengths of 488 and 543 nm, respectively. Live and dead cells were manually counted with QuPath[8], and the cell viability was calculated as the ratio of live cells to total cells.

### Supplementary Results

### S2.1. Quantitative assessment of *in vivo* $T_2$ mapping

In this study, *in vivo* $T_2$ mapping was conducted using a multi-echo spin-echo (MESE) sequence. MESE signals exhibit nonexponential decay patterns due to stimulated echoes and imperfect slice profiles[9]. To correct for these effects, the MESE signals were fitted to a simulated dictionary of MESE signals, which was created using the extended phase graph (EPG) method[7]. This dictionary included 360,000 MESE signal curves, spanning $T_2$ ranges from 20 to 200 ms and $B_1$ ranges from 0.5 to 1.5.

Supplementary Figure 1 illustrates the procedure to assessing the SNR and fitting quality for $T_2$ mapping using an example MESE image from the *in vivo* rat experiment. First, the background noise of MESE image was calculated. When a complex MR image contains Gaussian noise with a standard deviation of $\sigma$, the resulting magnitude image will exhibit Rician noise. This can be approximated as

Gaussian noise if the SNR (= $A/\sigma$) greater than 3, where $A$ is the magnitude of signal in the absence of noise[10]. Rician noise was sampled from 64 voxels in the background of the magnitude image and the noise level $\sigma$ was calculated from the average of the sampled Rician noise $M$ using the formula[10]:

$$M = \sigma\sqrt{\frac{\pi}{2}} \tag{S8}$$

Subsequently, a mask was created by thresholding the magnitude image at SNR > 3. $T_2$ mapping was performed by fitting the MESE signal to the simulated dictionary across the masked region. The fit quality was assessed using the normalized root mean squared error (NRMSE) and adjusted $R^2$, calculated as follows:

$$\text{NRMSE} = \frac{\sum(y_i - f_i)^2}{\sum y_i^2} \tag{S9}$$

$$\text{Adjusted } R^2 = 1 - \frac{n-1}{n-p-1}\frac{\sum(y_i - f_i)^2}{\sum(y_i - \bar{y})^2} \tag{S10}$$

where $\bar{y}$ is the mean of the signals, $y_i$ is the $i^{\text{th}}$ signal, $f_i$ is the $i^{\text{th}}$ fitted value, $n$ is the sample size (number of echoes), and $p$ is the number of variables.

Finally, the magnitude signal in the ROI beneath the perfusion chamber (width = 1.8 mm, depth = 0.6mm) was analyzed. After averaging the magnitude signals over the ROI, it is confirmed that the signal is well above the noise floor ($3\sigma$). After performing $T_2$ fitting, the fit quality was examined with NRMSE and adjusted $R^2$.

This analysis was conducted across all seven *in vivo* rat models. Supplementary Table S4 presents the fitted $T_2$ and $B_1$ value, NRMSE and adjusted $R^2$ value, the signal intensity of the last echo in the ROI, and the noise level $\sigma$. Notably, the maximum NRMSE was 0.043 and the minimum adjusted $R^2$ was 0.995, indicating a high degree of alignment between the observed magnitude signals and the fitting curves.

#### S2.2. Membrane potential-induced changes in $T_1$: SH-SY5Y cells

The changes of $T_1$ in SH-SY5Y cells when the membrane potential ($V_m$) was modulated by varying [$K^+$] = 0.2–80 mM or adding [$Ba^{2+}$] = 10 mM is shown in Supplementary Figure S2–3. A linear mixed-effect model was employed to analyze the effect of changes in $V_m$ ($\Delta V_m$) on $T_1$. This model included $T_1$ as a dependent variable, $\Delta V_m$ and its interaction with a group variable (indicating whether $\Delta V_m$ is induced by [$Ba^{2+}$] or [$K^+$]) as fixed effects, and cell batch as a random effect. The analysis revealed no significant interaction term ($p = 0.188$), showing that changes in $T_1$ do not depend on the specific method of membrane potential modulation. The model yielded the following relationship:

$$T_1\ (\text{ms}) = 1.94 \times 10^3\ (1 + 0.00197\ \Delta V_m) \cdots\cdots\cdots \tag{S11}$$

The effect $\Delta V_m$ on $T_1$ was significant ($p < 0.0001$), indicating increase in $T_1$ during depolarization and decrease during hyperpolarization. This trend is the same with $T_2$ regarding membrane potential, while the sensitivity is 3.5 times smaller compared to $T_2$. Subsequent post-hoc analysis compared each experimental condition to the control using Dunnett's test to account for multiple comparison, showing significant changes in $T_1$ across all conditions tested ($p < 0.0001$).

#### S2.3. Membrane potential-induced changes in $T_1$: Jurkat cells

The changes in $T_1$ were also evaluated using a different cell line, the Jurkat, under the same experimental conditions applied to SH-SY5Y cells. Each experimental condition was compared to the control using Dunnett's test to account for multiple comparison (Supplementary Fig. S4). As the same with observation in SH-SY5Y cells, Jurkat cells showed positive changes in $T_1$ during hyperpolarization and negative changes during depolarization, indicating that the detectability of membrane potential changes by $T_1$ does not appear specific to cell type.

### Supplementary Figures

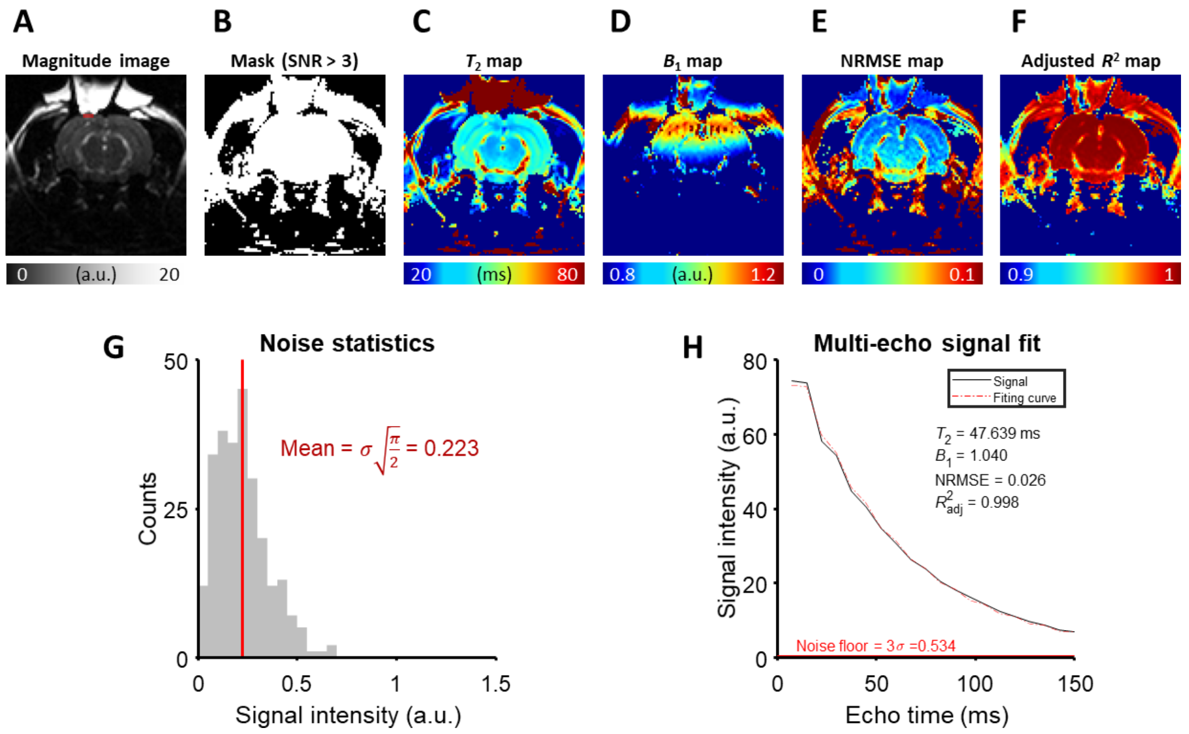


**Fig. S1**. The detailed analysis of $T_2$ fitting on an example MESE image from the *in vivo* rat experiment. (A) A magnitude image at the last echo time. The ROI is marked with a red rectangle. (B) A mask generated by thresholding the magnitude image with SNR > 3. (C) $T_2$ map, (D) $B_1$ map, (E) NRMSE map, and (F) Adjusted $R^2$ map produced by the fitting procedure. (G) The noise statistics of the background of the magnitude image. (H) The magnitude signal averaged over the ROI and its fitting curve.

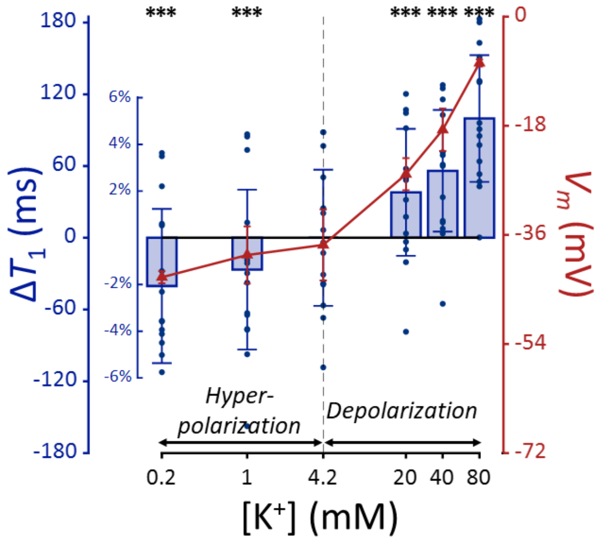


**Fig. S2.** $T_1$ and membrane potential ($V_m$) of SH-SY5Y cells versus extracellular $K^+$ concentrations ($[K^+]$). Changes in $T_1$ are displayed with blue bars ($n = 15$). Membrane potentials are plotted with red triangles ($n = 3$). The abscissa is in logarithmic scale. Error bars denote standard deviation. Statistical significance of changes in $T_1$ is marked with asterisks (***: $p < 0.001$).

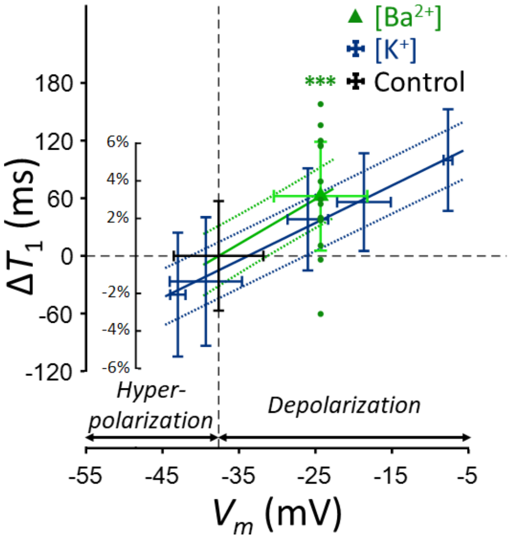


**Fig. S3.** Changes in $T_1$ of SH-SY5Y cells across experimental conditions: $[K^+] = 0.2$–$80$ mM (blue cross) and $[Ba^{2+}] = 10$ mM (green triangle), compared to the control condition (black cross). Data from fifteen experiments ($n = 15$) are displayed. Linear regression lines for $[K^+]$ data (blue solid line) and $[Ba^{2+}]$ data (green solid line) are drawn along with 95% confidence intervals. Error bars denote standard deviation. Statistical significance of changes in $T_1$ with $[Ba^{2+}] = 10$ mM is marked with asterisks (***: $p < 0.001$).

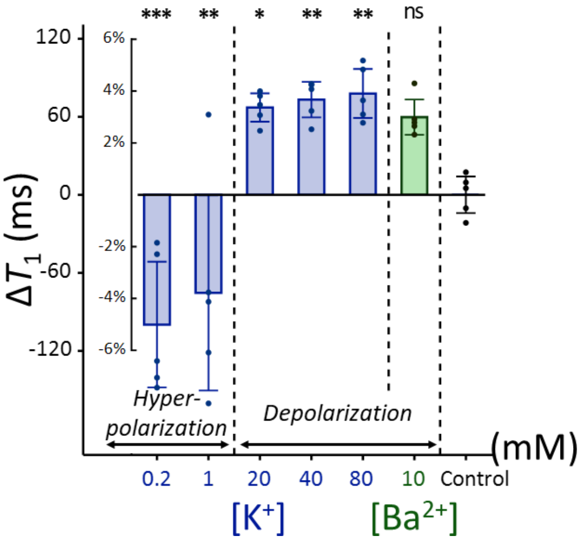


**Fig. S4.** Changes in $T_1$ of Jurkat cells across experimental conditions: $[K^+]$ = 0.2–80 mM (blue bar) and $[Ba^{2+}]$ = 10 mM (green bar), compared to the control condition of $[K^+]$ = 4.2 mM ($n = 5$). Error bars denote standard deviation. Statistical significance of changes in $T_1$ is marked with asterisks (ns: $p > 0.05$, *: $p < 0.05$, **: $p < 0.01$, ***: $p < 0.001$).

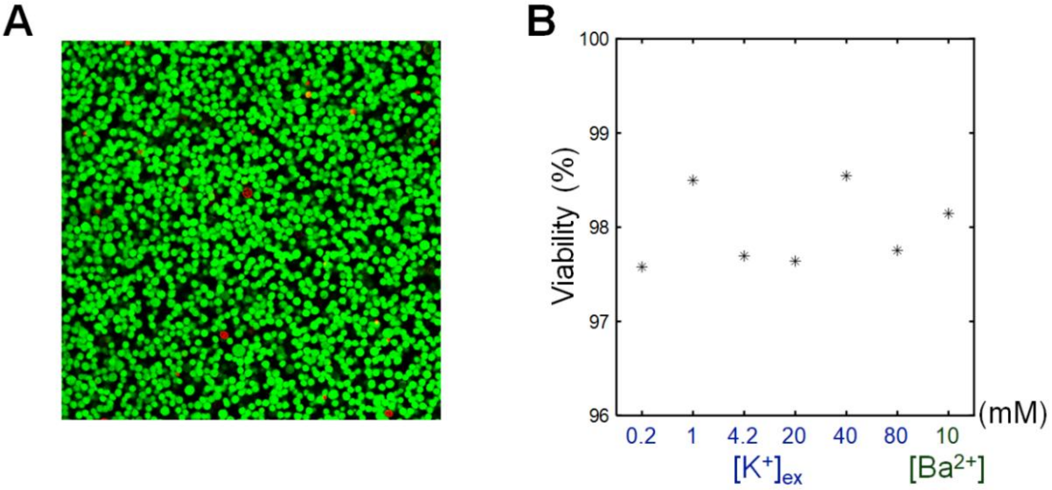


**Fig. S5.** The viability assay of SH-SY5Y cells. (A) A representative confocal microscopy image of an SH-SY5Y pellet. $[K^+]$ of extracellular medium was 4.2 mM. Live cells (green) were stained with calcein-AM, and dead cells (red) were stained with EthD-1. (B) The viabilities of SH-SY5Y cells versus the extracellular $K^+$ concentrations ($[K^+]$) and $Ba^{2+}$ concentration ($[Ba^{2+}]$).

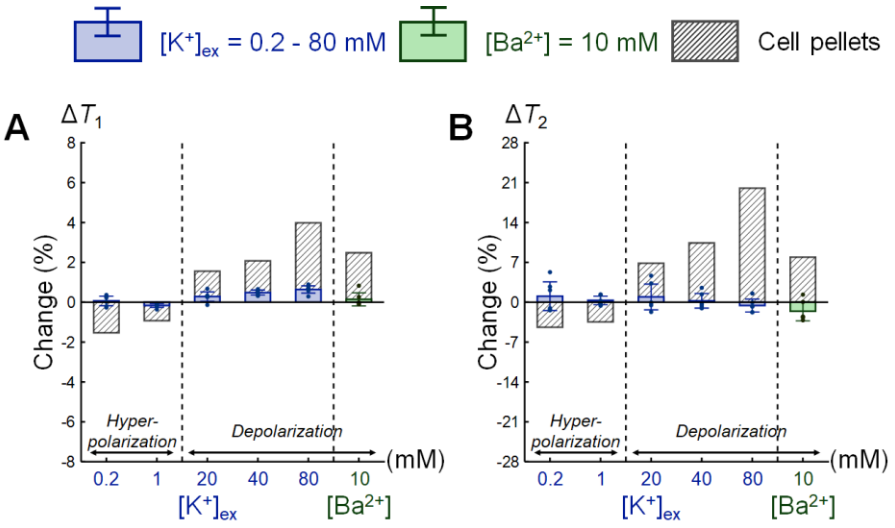


**Fig. S6.** Comparison of changes in relaxation times between extracellular media and SH-SY5Y cell pellets. Changes in (A) $T_1$ and (B) $T_2$ are expressed as percentage changes relative to the control condition of $[K^+] = 4.2$ mM. The results for extracellular media are displayed as blue bars for the extracellular $K^+$ concentrations ($[K^+]$) and green bars for the $Ba^{2+}$ concentration ($[Ba^{2+}]$) ($n = 6$). For comparison, the results for SH-SY5Y cell pellets corresponding to each extracellular medium are plotted. Error bars denote standard deviation.

### Supplementary Tables

**Table S1.** The scan parameters of the sequences used in the *in vitro* MR experiments. The cell samples were scanned with two types of sequences, single-echo spin-echo (SESE) and inversion recovery multi-echo spin-echo (IR-MESE) sequences, on a 9.4 T MRI. $T_2$ was estimated from the SESE sequence. $T_1$, PSR, and $k_{mf}$ were estimated from the IR-MESE sequence, and its inversion times (TIs) were optimized using the theory of Cramér-Rao low bounds: TIs = 4, 4, 4, 4, 4, 4, 4, 4, 17.91, 18.18, 18.18, 18.2, 18.2, 18.21, 18.21, 18.24, 18.24, 18.24, 18.24, 18.24, 18.31, 18.31, 18.32, 18.46, 55.18, 55.26, 55.53, 55.98, 163.5, 164.83, 164.92, 164.93, 164.96, 165.25, 165.25, 165.62, 197.68, 1976.77, 2280.95, and 10076.4 ms. The recovery time or repetition time was set long enough to ensure full relaxation of nuclear magnetization.

| **Sequence type** | SESE | IR-MESE |
|---|---|---|
| **Recovery time (ms)** | N/A | 15000 |
| **Repetition time (ms)** | 15000 | N/A |
| **Inversion time (ms)** | N/A | 4 to 10,079.4 (40 steps) |
| **Echo time (ms)** | 9.5 to 290.5 (linear, 50 steps) | 9.5 to 152 (linear, 16 steps) |
| **Resolution (mm)** | 0.5 | |
| **Slice thickness (mm)** | 1 | |
| **Estimated parameters** | $T_2$ | $T_1$, PSR, and $k_{mf}$ |

**Table S2.** The scan parameters of the sequence used in the *in vivo* rat MR experiments. The rats were scanned with a multi-echo spin-echo (MESE) sequence on a 7 T MRI.

| **Sequence type** | MESE |
|---|---|
| **Repetition time (ms)** | 1000 |
| **Inversion time (ms)** | N/A |
| **Echo time (ms)** | 7.5 to 150 (linear, 20 steps) |
| **Resolution ($mm^2$)** | 0.3 × 0.3 |
| **FOV ($mm^2$)** | 28.8 × 28.8 |
| **Slice thickness (mm)** | 2 |
| **Estimated parameters** | $T_2$ |

**Table S3.** The composition of the extracellular media used to modulate membrane potential *in vitro*. The sodium chloride concentrations were adjusted to maintain the same osmolarity across all media. Besides the inorganic salts listed in this table, all media commonly contained HEPES = 20 mM; glucose = 4.5 g/l; EGTA = 10 μM; pH = 7.2.

| **Medium type** | KCl (mM) | $BaCl_2$ (mM) | NaCl (mM) |
|---|---|---|---|
| **Baseline** | 4.2 | | 145.8 |
| **Low [$K^+$]** | 0.2 | | 149.8 |
| | 1 | | 149 |
| **High [$K^+$]** | 20 | | 130 |
| | 40 | | 110 |
| | 80 | | 70 |
| **[$Ba^{2+}$]** | 4.2 | 10 | 130.8 |

**Table S4**. The statistics of *in vivo* rat $T_2$ mapping experiments.

| Rat number | Condition | [$K^+$] (mM) | $T_2$ (ms) | $B_1$ | NRMSE | Adjusted $R^2$ | Noise level | Signal intensity at the last echo |
|---|---|---|---|---|---|---|---|---|
| 1 | Baseline | 3 | 47.6 | 1.04 | 0.026 | 0.998 | 0.18 | 6.98 |
| | Depolarization | 40 | 48.3 | | 0.025 | 0.999 | | 7.05 |
| | Depolarization | 80 | 48.8 | | 0.024 | 0.999 | | 7.21 |
| | Recovery | 3 | 47.9 | | 0.028 | 0.998 | | 7.12 |
| 2 | Baseline | 3 | 50.2 | 0.88 | 0.023 | 0.999 | 0.15 | 7.70 |
| | Depolarization | 40 | 50.7 | | 0.027 | 0.998 | | 7.76 |
| | Depolarization | 80 | 51.3 | | 0.027 | 0.998 | | 7.86 |
| | Recovery | 3 | 50.5 | | 0.032 | 0.997 | | 7.96 |
| 3 | Baseline | 3 | 45.4 | 0.85 | 0.033 | 0.997 | 0.19 | 3.19 |
| | Depolarization | 40 | 46.4 | | 0.028 | 0.998 | | 3.27 |
| | Depolarization | 80 | 46.5 | | 0.032 | 0.997 | | 3.33 |

| | | | | | | | | |
|---|---|---|---|---|---|---|---|---|
| 4 | Baseline | 3 | 42.9 | 0.79 | 0.019 | 0.999 | 0.16 | 5.47 |
| | Depolarization | 40 | 43.0 | | 0.023 | 0.999 | | 5.50 |
| | Depolarization | 80 | 43.4 | | 0.018 | 0.999 | | 5.60 |
| 5 | Baseline | 3 | 43.4 | 0.71 | 0.018 | 0.999 | 0.22 | 5.13 |
| | Depolarization | 40 | 43.7 | | 0.020 | 0.999 | | 5.33 |
| | Depolarization | 80 | 44.0 | | 0.017 | 0.999 | | 5.18 |
| 6 | Baseline | 3 | 55.1 | 0.78 | 0.037 | 0.995 | 0.35 | 6.32 |
| | Depolarization | 40 | 56.6 | | 0.033 | 0.996 | | 6.34 |
| | Depolarization | 80 | 56.9 | | 0.033 | 0.996 | | 6.90 |
| 7 | Baseline | 3 | 43.5 | 0.76 | 0.038 | 0.996 | 0.20 | 6.14 |
| | Depolarization | 40 | 44.2 | | 0.041 | 0.995 | | 6.25 |
| | Depolarization | 80 | 44.9 | | 0.039 | 0.996 | | 6.67 |

**Table S5.** The $T_2$ (ms) values of *in vitro* SH-SY5Y and Jurkat pellets.

| Pellet | $[K^+]$ (mM) | | | | | | $[Ba^{2+}]$ (mM) |
|---|---|---|---|---|---|---|---|
| | 0.2 | 1 | 4.2 | 20 | 40 | 80 | 10 |
| SH-SY5Y 1 | 46.77 | 47.26 | 48.53 | 52.68 | 54.57 | 58.86 | 52.35 |
| SH-SY5Y 2 | 47.41 | 47.53 | 48.45 | 51.92 | 53.3 | 58.07 | 52.22 |
| SH-SY5Y 3 | 46.19 | 47.12 | 49.76 | 54.45 | 56.24 | 60.59 | 54.98 |
| SH-SY5Y 4 | 46.69 | 47.08 | 49.11 | 52.29 | 54.85 | 57.57 | 52.64 |
| SH-SY5Y 5 | 49.04 | 49.01 | 51.64 | 54.16 | 55.38 | 63.76 | 56.53 |
| SH-SY5Y 6 | 47.4 | 48.63 | 50.19 | 52.85 | 54.23 | 59.6 | 53.82 |
| SH-SY5Y 7 | 45.54 | 46.45 | 48.85 | 52.07 | 55.26 | 57.75 | 51.42 |
| SH-SY5Y 8 | 45.21 | 46.17 | 47.59 | 52.38 | 53.58 | 57.7 | 53.58 |
| SH-SY5Y 9 | 48.69 | 48.08 | 50.88 | 53.7 | 56.18 | 61.24 | 56.24 |
| SH-SY5Y 10 | 46.18 | 47.08 | 49.3 | 53.25 | 54.64 | 59.63 | 55.49 |
| SH-SY5Y 11 | 47.62 | 47.66 | 49.36 | 53.3 | 54.64 | 59.95 | 56.18 |
| SH-SY5Y 12 | 52.01 | 51.63 | 52.12 | 56.63 | 57 | 64.85 | 55.63 |
| SH-SY5Y 13 | 50.85 | 51.62 | 54.23 | 58.63 | 60.68 | 62.59 | 55.39 |
| SH-SY5Y 14 | 50.51 | 52.47 | 52.69 | 54.45 | 57.11 | 62.21 | 55.06 |
| SH-SY5Y 15 | 50.66 | 50.14 | 51.25 | 52.43 | 54.45 | 60.42 | 51.38 |
| Jurkat 1 | 43.51 | 43.86 | 53.41 | 64.77 | 66.5 | 69.33 | 63.59 |
| Jurkat 2 | 41.37 | 38.5 | 54.33 | 62.48 | 63.46 | 63.9 | 59.46 |
| Jurkat 3 | 43.35 | 40.81 | 54.78 | 64.25 | 65.28 | 65.78 | 63.56 |
| Jurkat 4 | 48.08 | 44.82 | 55.59 | 59.94 | 61.97 | 61.19 | 59.64 |
| Jurkat 5 | 48.39 | 44.09 | 56.28 | 61.08 | 60.21 | 60.45 | 59.63 |

**Table S6.** The PSR values of *in vitro* SH-SY5Y and Jurkat pellets.

| Pellet | $[K^+]$ (mM) | | | | | | $[Ba^{2+}]$ (mM) |
|---|---|---|---|---|---|---|---|
| | 0.2 | 1 | 4.2 | 20 | 40 | 80 | 10 |
| SH-SY5Y 1 | 0.0395 | 0.0413 | 0.0388 | 0.0359 | 0.0366 | 0.0335 | 0.0347 |
| SH-SY5Y 2 | 0.0408 | 0.04 | 0.0365 | 0.0363 | 0.0345 | 0.0324 | 0.0367 |

| | | | | | | | |
|---|---|---|---|---|---|---|---|
| SH-SY5Y 3 | 0.0421 | 0.0382 | 0.0379 | 0.035 | 0.0345 | 0.0311 | 0.0364 |
| SH-SY5Y 4 | 0.0401 | 0.0385 | 0.0382 | 0.0344 | 0.0362 | 0.0337 | 0.0358 |
| SH-SY5Y 5 | 0.0359 | 0.0372 | 0.0361 | 0.0355 | 0.0342 | 0.0301 | 0.0307 |
| SH-SY5Y 6 | 0.0416 | 0.0381 | 0.0366 | 0.0345 | 0.0323 | 0.031 | 0.0324 |
| SH-SY5Y 7 | 0.0387 | 0.041 | 0.034 | 0.0347 | 0.0275 | 0.0328 | 0.034 |
| SH-SY5Y 8 | 0.0415 | 0.0412 | 0.0385 | 0.035 | 0.0346 | 0.0335 | 0.0357 |
| SH-SY5Y 9 | 0.0448 | 0.0457 | 0.0432 | 0.041 | 0.0393 | 0.0381 | 0.0384 |
| SH-SY5Y 10 | 0.0443 | 0.042 | 0.0405 | 0.0394 | 0.036 | 0.0365 | 0.0354 |
| SH-SY5Y 11 | 0.041 | 0.0418 | 0.0405 | 0.0369 | 0.035 | 0.0337 | 0.0353 |
| SH-SY5Y 12 | 0.0337 | 0.0324 | 0.0308 | 0.0308 | 0.028 | 0.0292 | 0.0293 |
| SH-SY5Y 13 | 0.0355 | 0.0362 | 0.0316 | 0.0322 | 0.03 | 0.0281 | 0.0309 |
| SH-SY5Y 14 | 0.0341 | 0.033 | 0.0374 | 0.0328 | 0.0306 | 0.0299 | 0.0298 |
| SH-SY5Y 15 | 0.0334 | 0.0354 | 0.0319 | 0.0336 | 0.0306 | 0.0272 | 0.0323 |
| Jurkat 1 | 0.0197 | 0.0152 | 0.0144 | 0.0145 | 0.0126 | 0.0116 | 0.0106 |
| Jurkat 2 | 0.0211 | 0.0225 | 0.0179 | 0.0131 | 0.013 | 0.0125 | 0.0148 |
| Jurkat 3 | 0.0177 | 0.0187 | 0.0162 | 0.0124 | 0.0106 | 0.011 | 0.0127 |
| Jurkat 4 | 0.0181 | 0.0155 | 0.0153 | 0.0143 | 0.0125 | 0.0124 | 0.0146 |
| Jurkat 5 | 0.0168 | 0.0174 | 0.0156 | 0.0122 | 0.0159 | 0.0146 | 0.016 |

**Table S7.** The $k_{mf}$ (Hz) values of *in vitro* SH-SY5Y and Jurkat pellets.

| Pellet | $[K^+]$ (mM) | | | | | | $[Ba^{2+}]$ (mM) |
|---|---|---|---|---|---|---|---|
| | 0.2 | 1 | 4.2 | 20 | 40 | 80 | 10 |
| SH-SY5Y 1 | 14.5 | 14.67 | 14.18 | 15.07 | 13.04 | 16.43 | 14.12 |
| SH-SY5Y 2 | 14.77 | 14.26 | 14.08 | 15.81 | 15.46 | 15.21 | 13.21 |
| SH-SY5Y 3 | 12.68 | 15.81 | 16.18 | 15.82 | 15.35 | 16.03 | 12.99 |
| SH-SY5Y 4 | 12.74 | 13.98 | 13.15 | 14.53 | 12.79 | 14.42 | 13.45 |
| SH-SY5Y 5 | 16.84 | 15.16 | 14.15 | 14.39 | 14.48 | 15.03 | 13.37 |
| SH-SY5Y 6 | 14.49 | 15.88 | 14.51 | 17.72 | 16 | 17.49 | 17.15 |
| SH-SY5Y 7 | 14.7 | 13.91 | 16.02 | 13.6 | 14.28 | 12.47 | 10.64 |
| SH-SY5Y 8 | 13.69 | 14.78 | 16.92 | 15.71 | 15.21 | 15.12 | 12.83 |
| SH-SY5Y 9 | 17.81 | 19.15 | 15.19 | 17.07 | 18.96 | 17.54 | 19.38 |
| SH-SY5Y 10 | 16.13 | 18.19 | 15.66 | 16.88 | 15.94 | 15.87 | 17.01 |
| SH-SY5Y 11 | 15.91 | 15.92 | 15.91 | 15.8 | 15.02 | 16.21 | 14.82 |
| SH-SY5Y 12 | 12.15 | 13.18 | 13.35 | 9.86 | 12.77 | 9.44 | 11.01 |
| SH-SY5Y 13 | 13.07 | 11.82 | 15.38 | 15.78 | 15.16 | 13.57 | 13.67 |
| SH-SY5Y 14 | 13.83 | 13.57 | 10.77 | 12.11 | 16.33 | 14.23 | 13.71 |
| SH-SY5Y 15 | 16.1 | 12.34 | 17.16 | 12.72 | 15.9 | 16.21 | 12.77 |
| Jurkat 1 | 21.17 | 11.31 | 13.39 | 19.54 | 11.36 | 12.35 | 20.15 |
| Jurkat 2 | 17.15 | 10.66 | 16.63 | 19.98 | 20.45 | 11.63 | 25.89 |
| Jurkat 3 | 12.78 | 15.25 | 17.29 | 15.79 | 24.83 | 25.39 | 25 |
| Jurkat 4 | 11.97 | 6.09 | 19.72 | 16.69 | 15.22 | 18.92 | 20.72 |
| Jurkat 5 | 18.52 | 25.26 | 15.25 | 14.25 | 10.98 | 19.62 | 15.97 |

**Table S8.** The $T_1$ (ms) values of *in vitro* SH-SY5Y and Jurkat pellets.

| Pellet | $[K^+]$ | | | | | | $[Ba^{2+}]$ |
|---|---|---|---|---|---|---|---|

| | (mM) | | | | | | (mM) |
|---|---|---|---|---|---|---|---|
| | 0.2 | 1 | 4.2 | 20 | 40 | 80 | 10 |
| SH-SY5Y 1 | 1865 | 1888 | 1896 | 1949 | 1956 | 2006 | 1949 |
| SH-SY5Y 2 | 1883 | 1915 | 1922 | 1956 | 1960 | 2000 | 1964 |
| SH-SY5Y 3 | 1872 | 1923 | 1949 | 2003 | 2014 | 2049 | 2011 |
| SH-SY5Y 4 | 1925 | 1932 | 1959 | 2003 | 2013 | 2044 | 1993 |
| SH-SY5Y 5 | 1965 | 1938 | 1973 | 2011 | 2024 | 2104 | 2067 |
| SH-SY5Y 6 | 1908 | 1934 | 1978 | 2001 | 2022 | 2084 | 2030 |
| SH-SY5Y 7 | 1883 | 1877 | 1940 | 1985 | 2069 | 2038 | 1992 |
| SH-SY5Y 8 | 1855 | 1890 | 1914 | 1970 | 1987 | 2030 | 1990 |
| SH-SY5Y 9 | 1811 | 1795 | 1844 | 1874 | 1898 | 1953 | 1892 |
| SH-SY5Y 10 | 1840 | 1855 | 1886 | 1932 | 1966 | 1996 | 1990 |
| SH-SY5Y 11 | 1876 | 1876 | 1915 | 1943 | 1967 | 2017 | 2007 |
| SH-SY5Y 12 | 1996 | 2039 | 2041 | 2057 | 2056 | 2116 | 2073 |
| SH-SY5Y 13 | 1963 | 1966 | 2003 | 2044 | 2043 | 2083 | 2068 |
| SH-SY5Y 14 | 2024 | 2038 | 2041 | 2073 | 2081 | 2136 | 2111 |
| SH-SY5Y 15 | 2021 | 2026 | 2030 | 2060 | 2078 | 2133 | 2089 |
| Jurkat 1 | 1866 | 1918 | 1983 | 2069 | 2079 | 2097 | 2049 |
| Jurkat 2 | 1853 | 1872 | 2003 | 2073 | 2078 | 2090 | 2040 |
| Jurkat 3 | 1845 | 1833 | 1972 | 2043 | 2044 | 2055 | 2046 |
| Jurkat 4 | 1957 | 2055 | 1999 | 2055 | 2058 | 2049 | 2052 |
| Jurkat 5 | 1948 | 1911 | 2011 | 2063 | 2075 | 2066 | 2079 |
